# Reduction of visual stimulus artifacts using a spherical tank for small, aquatic animals

**DOI:** 10.1101/2020.09.22.309419

**Authors:** Kun Wang, Burkhard Arrenberg, Julian Hinz, Aristides B Arrenberg

**Affiliations:** Werner Reichardt Centre for Integrative Neuroscience, Institute for Neurobiology, University of Tübingen, D-72076 Tübingen, Germany; Graduate Training Centre for Neuroscience, University of Tübingen, D-72076 Tübingen, Germany; Prudenter Agas Hamburg, 22149 Hamburg, Germany; Friedrich Miescher Institute for Biomedical Research, 4058 Basel, Switzerland

**Keywords:** geometrical optics, light reflection, refraction, dispersion, meniscus, receptive fields, optic tectum, zebrafish, glass bulb, computer graphics simulation, calcium imaging

## Abstract

Delivering appropriate stimuli remains a challenge in vision research, particularly for aquatic animals such as zebrafish. Due to the shape of the water tank and the associated optical paths of light rays, the stimulus can be subject to unwanted refraction or reflection artifacts, which may spoil the experiment and result in wrong conclusions. Here, we employ computer graphics simulations and calcium imaging in the zebrafish optic tectum to show, how a spherical glass container optically outperforms many previously used water containers, including Petri dish lids. We demonstrate that aquatic vision experiments suffering from total internal reflection artifacts at the water surface or at the flat container bottom may result in the erroneous detection of visual neurons with bipartite receptive fields and in the apparent absence of neurons selective for vertical motion. Our results and demonstrations will help aquatic vision neuroscientists on optimizing their stimulation setups.

## Introduction

In vision research, an important standard protocol is to present visual stimuli to immobilized animals while recording behaviors and/or neuronal activity ^1-6^. The visual system extracts features from stimulus patterns, and, accordingly, visual neurons can respond to a range of features such as contrast, motion direction, spatial frequency, sizes, locations and shapes of the visual stimulus ^5,7-11^. Therefore, presenting high quality visual stimulus patterns to the animal eyes is crucial for the investigation of visual functions and neural encoding ^12^.

A conventional mounting platform, equipped with a standard-size monitor or a small LED screen, is suitable for neural direction selectivity and receptive field (RF) mapping analysis of the vertebrate optic tectum ^7,13^, since the RFs of most tectal neurons are small ^13-15^ and visual stimulus parameters, such as contrast and spatial frequency, are easy to control with programmable hardware and software ^16,17^.

Due to the large RFs of some of the neurons as well as the large binocular fields of view in the lateral-eyed zebrafish ^18,19^, the stimulus delivery system needs to cover a large proportion of the visual space surrounding both eyes. Ideally the stimulus surface should completely cover their visual fields, and the platform supporting the animal should allow for a large accessible view field for the animals ^20-22^.

Furthermore, research using aquatic model animals (e.g. fish) gives rise to additional challenges, since an underwater environment is required during the experiments ^20,23^. Electronics need to be kept outside of the water, and therefore visual stimuli are oftentimes blurred and distorted before they reach the animal eyes, which mainly results from light refraction and reflection and the associated geometrical optics at the air-container-water interfaces ^12,24^. For example, during the presentation of global optic flow, the motion consistency across the visual field may be disrupted. Alternatively, the stimulus delivery system could be set underwater, surrounding the experimental animal. However, the waterproof protection for the electronic setup is usually difficult to ascertain and it may cause optical disturbances itself.

Here, we make use of Snell’s law, which posits that disruptive optical effects are quite small when light beams pass through an optical interface orthogonally, and design a new spherical glass bulb container, coupled with an adjustable rotation mount holder to optimize vision experiments in small-size fish. Moreover, we demonstrate the optical advantages of the new glass bulb using optical simulations and functional neural activity recordings.

## Results

### Design and optical advantages of a spherical glass container

The underwater presentation of visual stimuli can suffer from a range of optical artifacts, such as total internal reflection (TIR), light refraction, general reflection, dispersion, water meniscus, and light absorption (Figure 1e). Given the different refractive indices of water, air, and the container material, the shape of the container (e.g. flat vs. round walls) will have a profound impact on the path of the light transitioning into water. In many previous studies on zebrafish vision, the potential occurrence of stimulus artifacts had received little attention ^12^. Here, we compare the occurring optical artifacts of three different containers for aquatic animals: a commonly used Petri dish lid (Figure 1b), a cylindrical water container (Figure 1c), and a new spherical glass container that we designed to minimize artifacts (Figure 1d). The glass bulb container is 8 centimeters (cm) in diameter and has an opening of 4.6 cm (in diameter) on the top, which allows for *in vivo* microscopy using a water immersion objective from above (Figure 1d, Supplementary Figure S1). During the experiment, the animal is immobilized on the tip of a triangular stage located in the center of the spherical glass bulb (Supplementary Figure S1). In comparison to the Petri dish lid and the cylindrical container (Figure 1b, c and Supplementary Figure S1), the spherical glass bulb allows for larger homogeneously accessible visual space (see the spherical panorama views covering 360° in azimuth and 180° in elevation in Figure 1f-h). In addition, using a rotation mount metal holder (Supplementary Figure S1), the animal’s position is adjustable around the three axes of the Cartesian space.

**Figure 1.**
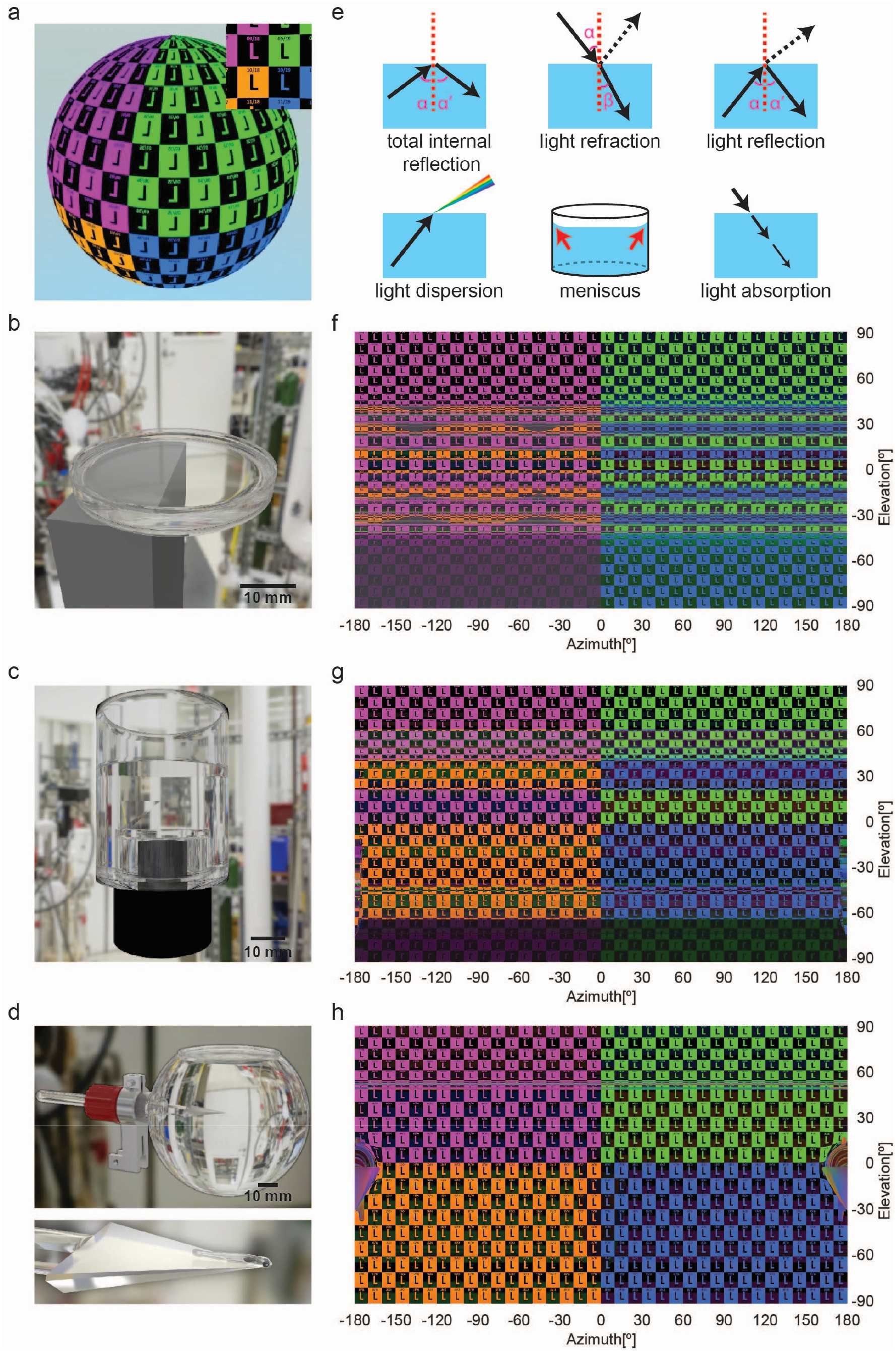
Simulation of visible stimulus patterns in three different water containers. (**a**) The sphere shows the simulated checkerboard stimulus pattern (18 rows, 36 columns) under ideal optical condition. The letter “L” was used to keep track of the orientation (up, down, left, and right) of stimulus patches. Inset: a patch of the checkerboard stimulus (0° in elevation; 0° in azimuth) seen from the inner center of the container. (**b-d**) Photorealistic illustrations of three water containers (3D rendering). (**b**) A commercially available Petri dish lid (Ø 35 mm) on a metal stage. (**c**) A plastic cylindrical container made by a fine mechanics workshop (Ø 40 mm) on a camera lens holder. (**d**) A custom-made glass bulb (Ø 80 mm) attached to a holder. Below, a simulated larval zebrafish is on the tip of a triangular stage. (**e**) Illustration of optical effects: total internal reflection, light refraction, reflection, dispersion, water meniscus, and light absorption. Black arrows represent light beams and vertical red dashed lines indicate the perpendiculars. Angles α, α’, and β indicate angles of incidence, reflection and refraction. Red arrows indicate the water meniscus at the container wall. (**f-h**) 360° panorama picture of the simulated visual patterns seen from the perspective of the animal in the center of the corresponding containers. (**f**) Petri dish lid, (**g**) cylindrical container, and (**h**) glass bulb.

### Simulation of optical stimulus artifacts in different containers

We simulated the details of the visual stimulus patterns for the three water containers as perceived by the fish’s eyes (Figure 1f-h). In the simulation, the upper visual space was covered by two quarter spheres of purple and green colored checkerboard stimuli. The lower visual space was covered by orange and blue stimuli (Figure 1a).

Strong artifacts are present for Petri dish lids around the equator (−41.4° to +41.4° in elevation, Figure 1f). For example, in our simulation the animal’s right eye sees a pattern of alternating green and blue stimuli, so stimuli from the upper and lower visual space are inappropriately mixed. These stimulus artifacts result from total internal reflection (TIR) and are also visible to animals placed inside the cylindrical container (Figure 1g). Any light ray that penetrates into the water through the air-plastic-water interface of Petri dish side wall with an incidence angle smaller than 62° (Supplementary Figure S2) is subject to TIR. Furthermore, the visual space above the water surface (and at the flat Petri dish bottom) is compressed into visual field elevations ranging between roughly 41.4° and 90° (and roughly -41.4° to -90° for the Petri dish bottom), which corresponds to the so-called Snell’s window. Thus, in very flat water containers such as the Petri dish lid, visible stimuli in the elevation range around the equator (roughly - 40° to +40°) correspond to stimulus light rays that entered the water body through the side walls of the Petri dish. Many of these light rays are subject to multiple rounds of TIR reflection at the water surface and the Petri dish bottom before they reach the animal’s eye (Supplementary Figure S2). These TIR stimulus artifacts are absent in the spherical container (Figure 1h), since only the light beams with incidence angles close to 0° can reach the fish’s eyes located in the sphere center, thus preventing visible TIR (Supplementary Figure S5).

Furthermore, the stimulus pattern observed by the animal’s eye in the Petri dish lid gets distorted via refraction when the light beams intersect the air-to-water border, i.e. at the water surface, or through the bottom of the Petri dish lid. Strong refraction is observed for large incidence angles (α) of light (Figure 1e), which causes stimulus compression along the vertical axis at elevations around 50° (Figure 1f), while stimuli near the poles (close to +/-90° elevation) still have the squarish aspect ratio present in the undisturbed checkerboard stimulus. These refractive distortions are also present in the cylindrical container (Figure 1g, elevation range from 45° to 90°). In the spherical container, the opening for the microscope objective still causes refractive distortions for elevation angles exceeding 60° (Figure 1h). However, the upper part of the visual field would in any case be blocked by the microscope objective during experiments requiring microscopy (Supplementary Figure S1, elevation angles exceeding 41.6° are blocked).

In addition to TIR and refraction, general light reflection artifacts occur in all three containers on both the contralateral and ipsilateral sides of the presented visual stimuli. Due to the similar refractive indices of the containers, the reflectance on the contralateral side is quite similar for all three containers. The reflectance is relatively low, for small incidence angles of light (Petri dish lid, 5%; cylindrical container, 4%; glass bulb, 4%). Though weak, the contralateral reflection is still visible to the contralateral eye on a dark background (Figure S4c-d). The effects of the reflection can influence monocular experiments as shown experimentally further below.

Reflection on the ipsilateral side is very weak in case of the glass bulb and the cylindrical arena (0.14% and 0.16, respectively). In the Petri dish lid, however, the ipsilateral reflectance is high (first reflection approx. 25%, second reflection, approx. 15%;) when light rays come from above or below. For example, in the lower visual field (−40° to -90° elevation) of the left eye (0° to - 180° azimuth), a reflection of the upper visual space (purple-colored stimuli) is faintly visible (See Figure 1f). Furthermore, depending on the water level in the Petri dish lid, these lateral reflections can be partially blocked by the objective due to its small access angle. In contrast, the reflection resulting from the light rays directly entering from the side is negligible for the Petri dish lid as well (0.29%).

Light dispersion is restricted to non-orthogonal light beams passing through the water surface (Figure 1e and Supplementary Figure S1). Therefore, it does not occur in the glass bulb since only orthogonal light beams reach the visual field of the stimulated animal eye via the bulb. For the Petri dish and cylindrical container, light disperses on the non-spherical polystyrene surfaces. Blur and rainbow edges resulting from chromatic aberration are visible to the animal (Supplementary Figure S1). The light dispersion artifacts are, however, very weak in comparison to reflection and refraction artifacts, especially when working with narrow-bandwidth (monochromatic) LEDs (Supplementary Figure S1).

Modelling the surface of the water close to the edge of the containers (Figure 1e and Supplementary Figure S1) revealed that the water meniscus only interrupts the narrow range of upper elevations (e.g. 50° in elevation in the spherical container, Figure 1h and Supplementary Figure S1). Due to the relatively shallow access angles of microscope objectives (e.g. 41.6°, Supplementary Figure S1), the water meniscus and rim do not affect the upper visual field of the fish in the spherical and cylindrical containers. For the Petri dish lid, the meniscus artifacts are visible to the animal, but they are much less problematic than the above described light refraction and reflection artifacts (Figure 1f).

Next to directional changes of light rays, which were discussed above, a fraction of the light also gets absorbed in the water. Due to the small sizes of the containers, light absorption within pure water does not exceed 0.2% for green stimulus light ^25^ and, therefore, is negligible for our analysis. While the transparency of the three containers compared here is similar, edges or thickenings of the container (e.g., at the vent edges and the outer reinforcement ring of the Petri dish lid, or at the glued faces of the cylindrical tank increase opacity Figure 1b, c and Supplementary Figure S1).

The zebrafish has a large visual field of about 163° and the accessible visual space is even larger when taking eye movements into account ^26,27^. For each container in this study, its holder blocked parts of the stimulus (Figures 1f-h). Accordingly, a compact design of the holder is recommended to minimize the visible holder silhouette. For example, in our Petri dish lid simulation, the lid holder blocks the stimulus of the lower field of vision of the left eye (orange colors, Figure 1b, f). Due to the camera and bracket mounted at the bottom of the cylindrical container, the visual field downwards from -60° (in elevation) is blocked for all azimuth angles (Figure 1c, g). In contrast, for the spherical container, the mount holder, the glass rod (stage holder), and the wedge-shaped glass stage only occlude a small region of visual space in the rear of the animal (Figure 1d, h and Supplementary Figure S1).

In summary, the new glass bulb designed in this study offers a better optical environment to present visual stimuli to small aquatic animals than the other two containers. There is no disruption by TIR or light refraction and remaining reflection is very weak. Furthermore, stimuli covering a larger range of visual space can be presented using the glass bulb.

### Total internal reflection (TIR) disturbs receptive field mapping experiments in the optic tectum

When the water level is low, TIR occurs even in the spherical container and thus influences the visual perception of the fish. Stimuli presented at certain positions below the equator are reflected and visible to the animal in its upper visual field (Figure 2a), resulting in stimulus inconsistencies above the equator (Figure 2b). To demonstrate the effects of TIR on vision experiments, we mapped the receptive fields (RFs) of zebrafish tectal neurons using calcium imaging with a two-photon microscope ^8^.

**Figure 2.**
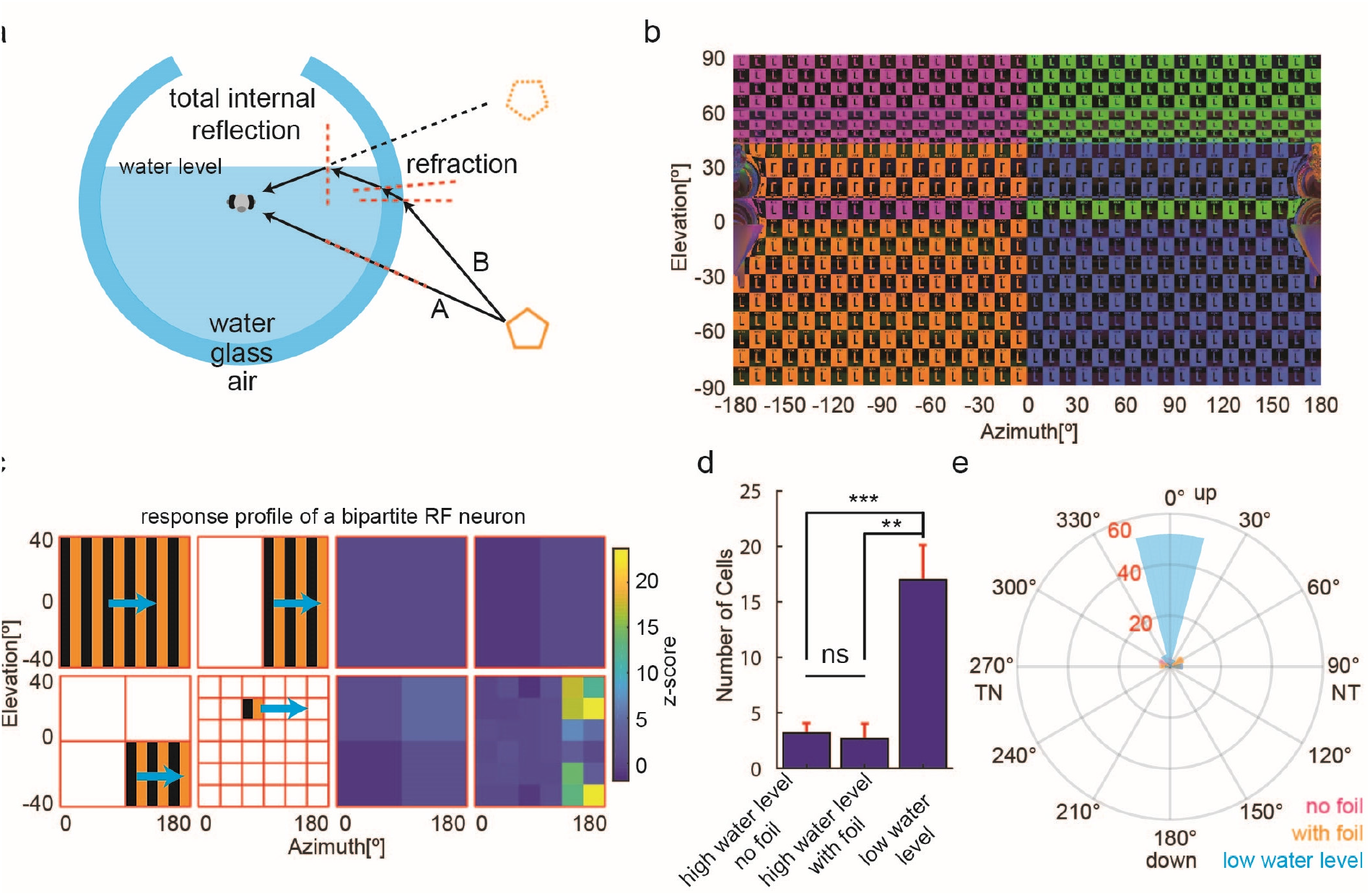
TIR results in the erroneous detection of bipartite receptive fields in the zebrafish optic tectum. (**a**) Illustration of TIR for the glass bulb with low water levels. Light beam A is perpendicular to the air-glass-water interface and reaches the fish right eye directly. However, light beam B, from the same light source as light beam A, is refracted twice on the air-glass-water interfaces and then reaches the fish’s eye from above via TIR. Black arrows, light beams; dashed red lines, perpendiculars; orange pentagon, one object; dashed orange pentagon, reflection of the object. (**b**) A 360° panorama picture of the stimulus pattern seen from the glass bulb center with low water level. Part of the visual stimulus below the equator is reflected to the upper visual field (the upside-down letters “L” in orange and blue) via TIR. (**c**) The response profile of a bipartite RF neuron. Left side: a diagram of the monocular RF mapping protocol (Supplementary Figure S3 shows the whole protocol). Horizontally moving gratings (indicated by cyan arrows) of different sizes and locations were sequentially presented to the right eye of the animal. Right side: z-scores of the calcium signal of a bipartite RF neuron responding to motion phases on the left. The neuron responded exclusively to small-size motion. Two separate RF centers are present in the same neurons and located in the upper and lower temporal visual fields, respectively. (**d**) The average numbers of bipartite RF neurons in each tested fish measured under different experimental conditions. ***, p<0.001; **, p<0.01; ns, no significant differences. The “high water level” group was further split into two subgroups (cf. Figure 3) for which the left eye was either occluded (“with foil”) or left free to see potential contralateral reflections (“no foil”) (**e**) A polar histogram of the orientation of the two RF centers for each neuron. Orientations of 0° and 180° correspond to a vertical orientation, 90° and 270° to a horizontal orientation of the bipartite RF centers. TN, temporal-nasal direction; NT, nasal-temporal direction. Neuron numbers are indicated in red. n = 6 (3 composite fish brains, high water level no foil), 6 (3 composite fish brains, high water level with foil) and 4 (2 composite fish brain, low water level) recordings in (d) and (e).

In the first condition, we filled a glass bulb (Ø 10 cm) up to 11.5° in elevation (1 cm water level above the animal, Figure 2a). In the second condition, we filled a glass bulb (Ø 8 cm) completely with water (up to ∼45° in elevation). We mapped visual RFs of larval zebrafish tectal neurons by using horizontally moving gratings of different sizes and locations that were presented to the right eye of the animal (Figure 2c and Supplementary Figure S3, see Methods). While most motion-sensitive tectal RFs should have a single, unimodal RF according to previous work ^8^, in the low water level condition (with visible TIR) some of the neurons (68 out of 543) appeared to have bipartite (double-field) RFs with two RF centers (Figure 2c, d). The two RF centers were vertically aligned for the vast majority of these bipartite RFs (Figure 2e). Therefore, this “bipartite RF” type is most likely a TIR artifact resulting from light reflection at the horizontal water surface (Figure 2a, b).

Tectal small-size RF centers cover nearly the whole monocular visual field in larval zebrafish and are biased to the upper nasal visual field (Supplementary Figure S3) ^8^. In this hotspot region, however, only very few small-size tectal RF were identified when the glass bulb was not filled up with water completely (Supplementary Figure S3). The small number of detected neurons was likely a combined result of TIR, reflection and water meniscus stimulus artifacts.

### General reflection disturbs monocular receptive field mapping experiments

As discussed above, reflections of the stimulus are visible for the animal on the contralateral side (Figure 3a-b). Any “visible” point of the stimulus display emits light rays that reach the ipsilateral eye (light beam a in Figure 3a), as well as light rays that miss the ipsilateral eye, part of which can be seen as reflection by the contralateral eye in the spherical container (light beam b in Figure 3a). The visible reflected light intensity is only about 4% of the original light (Figure 3c, d, Fresnel equations), since the visible light rays hit the air-glass-water interface nearly perpendicularly (e.g. light beam A in Supplementary Figure S5). The Petri dish and cylindrical containers suffer from similar reflections (Supplementary Figure S4.

**Figure 3.**
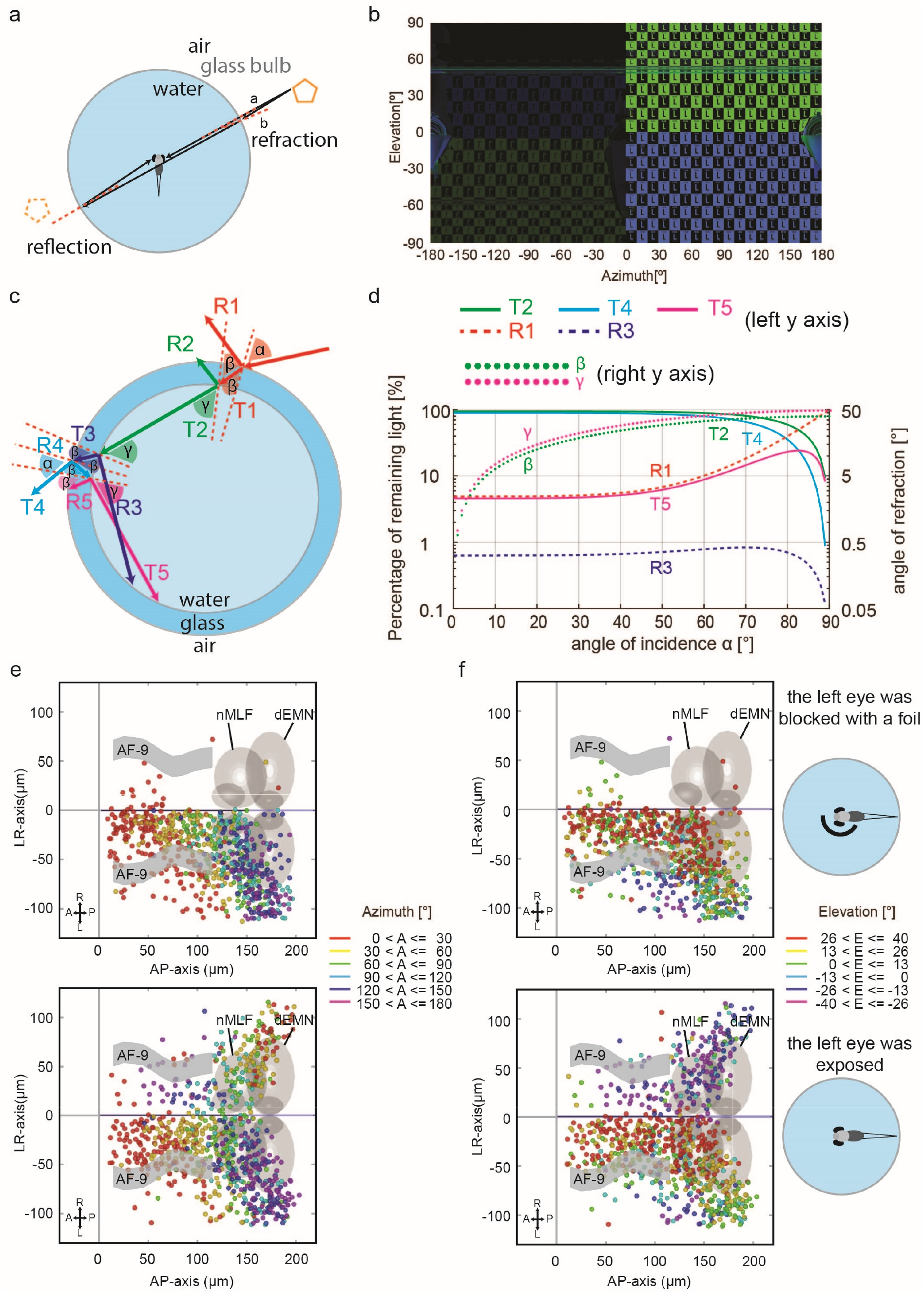
Remaining reflections in the glass bulb are visible to the animal. (**a**) Beam a, perpendicular to the air-glass-water interface, reaches the fish’s right eye directly and is absorbed. Light beam b, with a small but non-zero incidence angle at the air-glass-water interface, passes by the fish and is reflected back to the “non-stimulated” left eye. (**b**) 360° panorama picture of a half sphere stimulus: In the left hemifield (0° to -180° azimuth), the “non-stimulated” eye can see the point-symmetric reflection (left) of the monocular stimulus (right). (**c**) Detailed illustration of light reflection and refraction in the glass bulb. Angles α and β, angles of incidence and refraction on the air-glass interface; γ, the refraction angle at the glass-water interface. Since the glass is thin and the angle β is relatively small, the incidence angle at the glass-water interface approximates refractive angle β at the air-glass interface. Only light beams with relatively strong high power are shown here. Light beams R3 and T5 can reach the “non-stimulated” eye when the incidence angle α is small. ‘R’ and ‘T’, reflected and transmitted light beams. (**d**) Final reflection and refraction rates of the light, and angles of refraction calculated according to the Fresnel equation and Snell’s law. Some of the light beams in (c) have roughly equal light intensities and were therefore omitted in the plot in (d): T1=T2=T3; R2=R3=R5; R4=T5. (**e-f**) Topographic maps (in azimuth (e) and elevation (f)) of RF centers of small-size RF tectal neurons in the zebrafish (dorsal views). The “non-stimulated” (left) eye was covered in the first row (control group) but exposed to potential stimulus artifacts below (experimental animals). Each colored dot represents a single neuron with its receptive field center in the corresponding azimuth or elevation range. For example, neurons with RF centers between 0° (in front of the fish) and 30° azimuth on the nasal right side of the fish are in red in (e). n = 6 fish for both groups, 3 composite brains.

We tested the influence of the visible reflection to the “non-stimulated” eye using tectal RF mapping and a spherical container completely filled with water. In the control group, the non-stimulated eye (left eye) was blocked by a black foil. In these control animals, the active tectal neurons were mainly located in the tectal hemisphere contralateral to the stimulated eye as expected from the complete midline crossing of retinal ganglion cell axons in the optic chiasm. The RF centers furthermore showed the expected topographical distribution along the anterior-posterior, medial-lateral, and dorso-ventral axes (Figure 3e, f and Supplementary Figure S4). In contrast, in the experimental group without occlusion of the non-stimulated eye, many tectal neurons were identified in the ipsilateral hemisphere as well, and these ipsilateral neurons distributed in the tectum according to a reverse topographic map (Figure 3e, f and Supplementary Figure S4). These results suggest that the left eye (the “non-stimulated” eye) saw the weak stimulus reflection and that this reflection was strong enough to activate a small proportion of neurons in the corresponding tectal hemisphere. The roughly point-symmetric nature of this reflection also explains the occurrence of a topographic map with apparent reverse order (Figure 3e, f and Supplementary Figure S4).

### TIR and light refraction in the Petri dish lid result in inaccurate detection of preferred directions in zebrafish tectal neurons

The inconsistency and disruption of the visual stimulus patterns in the Petri dish lid are mainly caused by TIR at the horizontal interfaces. In vision experiments, this should lead to stimulus artifacts of different severity for vertical and horizontal motion directions. For shallow elevation angles in the Petri dish, each light ray bounces within the water before reaching the fish’s eye and the bounce number depends on the vertical angle of view. For even and odd numbers of reflection, the vertical moving directions of the visual stimuli projected to the fish’s eye are opposite. As a result, the vertical motion of simple grating bars is seen as motion in opposite vertical directions for odd numbers of TIRs (Figure 4a). Clearly, this should strongly reduce responsiveness of vertical direction-selective neurons that have a large receptive field. In contrast, such direction-inverting stimulus artifacts should be completely absent for horizontal motion directions of the stimulus, since these motion directions are parallel to the TIR-inducing water surface and lid bottom (Figure 4a).

**Figure 4.**
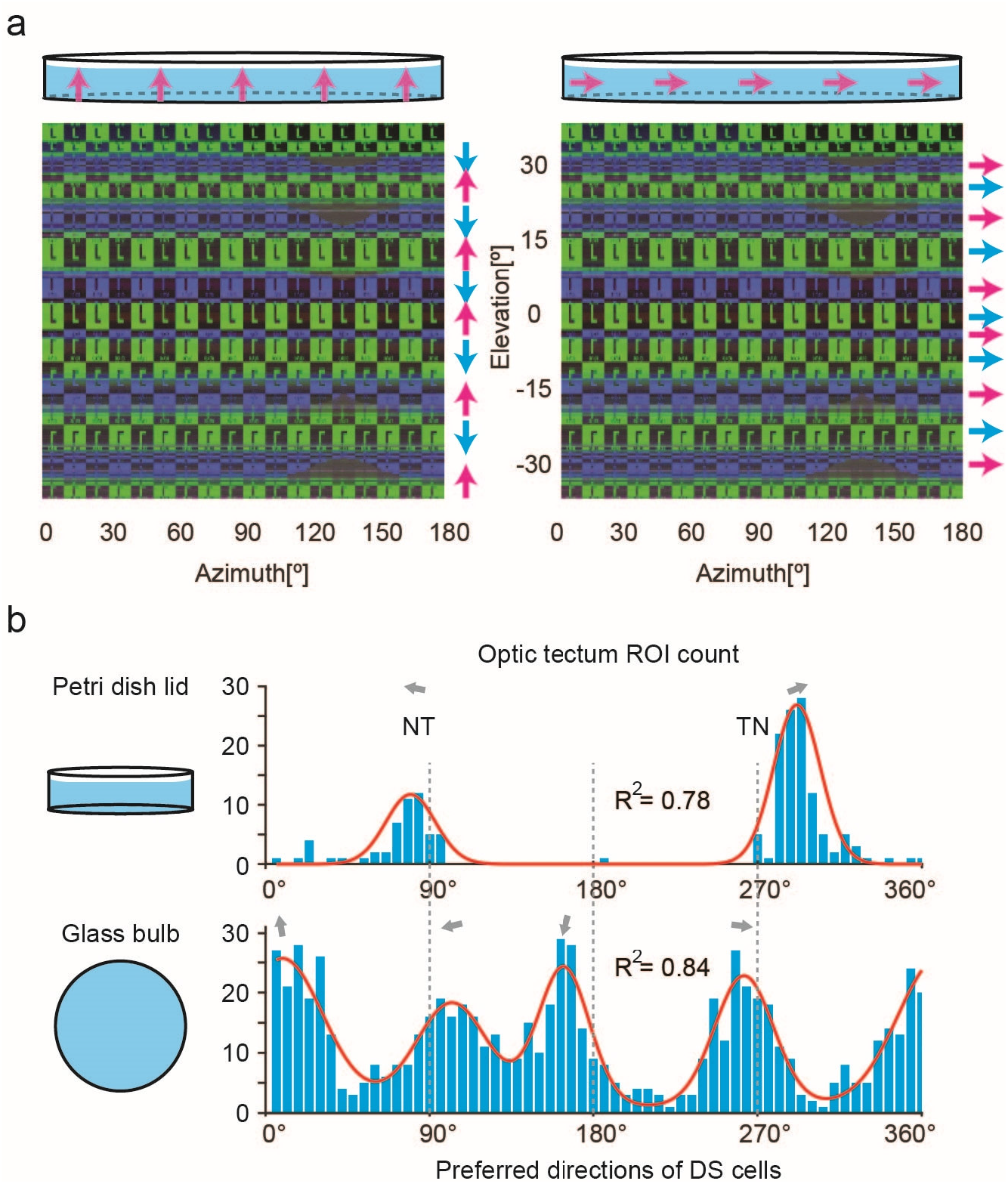
Vertical motion stimuli are disrupted due to TIR. (**a**) 180° panorama of the right side of the stimulus seen from the center of a Petri dish lid. Due to TIR, the visual stimulus patterns near the equator (−41° to 41° in elevation) are intermingled and oftentimes oriented upside-down (background colors and letters ‘L’, also see Figure 1f). Left side: vertical upward motion (indicated by magenta arrows) is presented from outside of a Petri dish. The whole moving pattern is disrupted and not continuous from the perspective of the animal in the center of the container. At different elevation levels of the visual field, the stimulus patches move upwards (magenta arrows) or in the opposite direction (cyan arrows). Right side: horizontal motion (indicated with magenta arrows) to the right is presented from outside of a Petri dish lid. Seen from the center of the Petri dish, the whole moving pattern is still continuous, consistent and homogeneous (cyan and magenta arrows). (**b**) Histograms of the preferred directions of direction-selective tectal neurons recorded in a Petri dish lid (top, n = 5 fish) or a glass bulb (bottom, n = 9 fish). The peaks were fitted with a sum of two (top) or four (bottom) Von-Mises functions (red lines). The PDs are indicated with gray arrows.

To demonstrate how direction selectivity analysis gets compromised by this artifact, we presented motion in eight different directions using a Petri dish lid (in this study) or a glass bulb as the container for the animal (Supplementary Figure S3) ^28^. Using the glass bulb, we appropriately detected direction-selective tectal neurons, which preferred either vertical or horizontal directions. For each of the four preferred directions (PD: up, down, nasalwards, and temporalwards), we found an approximately equal number of neurons in the tectum ^28^. In contrast, but as expected from our optical analysis above (Figure 4a), almost no direction-selective neurons preferring upward or downward stimulus motion could be detected in the experiments using the Petri dish lid. The histogram of preferred directions was instead dominated only by two peaks, nasalward and temporalward directions. The two PD peaks point a little upward compared to the corresponding horizontal PDs recorded with the glass bulb. The nasal-temporal direction (292°, 109 cells) is represented by many more neurons than the opposite temporal-nasal direction (80°, 44 cells; z-test for proportions, p<0.00001) (Figure 4b).

## Discussion

Our study identifies an alarming number of optical stimulus aberrations in commonly used containers for aquatic animals in vision research. We provide a solution in the form of a spherical glass container that can minimize most of the identified optical artifacts. Using calcium imaging, we also show the effects of these optical artifacts on RF mapping and direction selectivity analysis, thereby demonstrating the need for careful experimental design.

### The glass bulb offers a better optical presentation of the visual stimulus than the other two containers

Visual stimuli seen from within the Petri dish lid are severely disrupted across about 80° in elevation near the equator by TIR (Figure 1f). The bouncing light beams (Supplementary Figure S2) cause three caveats. First, visual stimuli from different spatial locations lead to stimulus overlap and blurring when they are projected to the same region within the fish’s eye. Second, stimuli from adjacent spatial locations project to angular regions far away in the eye (e.g., 30°), disrupting the stimulus pattern from the perspective of the animal. Third, at several elevation levels of the visual field, stimuli are mirrored by reflections (Figure 1f). For a vertical whole-field motion stimulus, this results in visible opposing vertical directions (Figure 4a) and a corresponding loss of direction-selective neuronal responses. Furthermore, light refraction leads to vertical stimulus pattern compression for certain elevation levels. Therefore, the Petri dish lid is clearly a suboptimal container for aquatic animal visual research, especially for experiments including presentation of vertical motion or small-size stimuli in specific locations used for RF mapping or prey capture experiments.

The cylindrical container offers a better optical environment than the Petri dish lid, though a relatively large part of the visual field is still disrupted by light refraction and reflection (Figure 1g). The cylindrical container should be designed to be as large as possible in the vertical direction (high on the top and deep at the bottom) to minimize the disturbance by its holder, water meniscus, light refraction and light reflection. Very large cylindrical containers require highly waterproof objectives for microscopy.

From the perspective of the animal in the container, the distortion and blur of the visual stimulus are dramatically minimized in a spherical glass container (Figure 1h), although several potential caveats still persist. First, it is important to place the animal in the center of the glass sphere, since mispositioning the fish in the horizontal plane away from the spherical center results in stimulus distortion near the two poles (Supplementary Figure S5). Second, the custom manufacture of glass bulbs with a perfect spherical curvature and homogeneous thickness can be difficult. Fortunately, the visual stimulus quality is quite robust against inaccurate manufacturing (Supplementary Figure S5). Third, our monocular RF mapping experiment revealed that the low reflectance (4% of the original intensity) at the glass-air interface is strong enough to activate tectal neurons. A further reduction of reflectance could potentially be achieved by optical coatings. Alternatively, the inner surface of the glass could be coated with a diffusive paint and used as back-projection screen for the stimuli. Occlusion of the non-stimulated eye is strongly recommended in monocular experiments with the glass bulb or with any water container.

### Influence of geometrical optics on RF mapping and direction selectivity analyses

Many apparently bipartite RF neurons were detected in the larval zebrafish tectum when the water level in the glass bulb was low and allowed TIR to be visible (Figure 2). However, not all of the small-size RF tectal neurons with RF centers above the equator responded to the reflection (Supplementary Figure S3). We speculate that the low intensity and contrast of the reflection - relative to those of the cardinal stimulus - prevented these neurons from being detected as bipartite neurons in our analysis.

In our investigation of direction-selective neurons, only two out of the four previously reported neuronal populations of direction-selective neurons were detectable when a Petri dish lid was used for the experiment (Figure 4b) ^28^. The incomplete detection most likely resulted from the disruption of the vertical motion by TIR (Figure 4a).

In summary, we demonstrate that optical artifacts can disturb key parameters such as motion directionality, and they can displace the position of the visual stimulus in aquatic vision experiments. Use of a spherical water container can greatly reduce artifacts. Our results showcase potential pitfalls in experimental design and provide a roadmap for careful design of vision experiments in aquatic environments.

## Methods

### 3D visualization techniques

All 3D visualizations were performed with the 3D computer graphics software Blender v2.79b (https://www.blender.org/download/releases/2-79/). The different experimental environments were modelled in detail and rendered in pictures to illustrate different effects. In this study, geometrical optical effects resulting from light transmission (absorption), reflection (including TIR), refraction, dispersion and occlusion, were calculated, simulated, and visualized.

#### Containers

We simulated three different fish containers based on popular usage in zebrafish vision research.

- Petri dish lids (Greiner 627102 with Ø 35 mm, Figure 1b) Zebrafish were embedded using agarose, at the bottom of the lids and the lids transferred to the platform for imaging ^29^.
- Cylindrical container (Figure 1c) Zebrafish were immobilized on a small, custom-made stage in the middle of the cylinder, with the low melting agarose.
- Glass bulb (Figure 1d) Zebrafish were immobilized on a small triangular stage, which were then placed in the center of the water filled glass bulb ^8^.

The different mounting devices were modeled together with their respective containers.

#### Inputs

The main inputs, indices of refraction (IOR) for different containers, are listed in the table below for the photorealistic modeling.

**Table 1.** The materials and the corresponding indices of refraction from the three containers.

| Containers | Petri dish lid | Cylindrical container | Glass bulb |
| --- | --- | --- | --- |
| Material | Polystyrene | Acrylic glass | Glass Schott Boro |
| IOR | 1.57 | 1.49 | 1.473 |

#### Stimulus

For optical simulations, a colored checkerboard pattern, consisted of 36 columns and 18 rows, was used with different colors to differentiate the main directions (up/down and right/left, marked by a letter ‘L’ to identify mirroring).

#### Renderer and cameras

The images were usually rendered using the Cycles renderer of Blender. For photorealistic representation we mostly used an f = 35 mm perspective camera (e.g., Figure 1b-d). For the test of the optical quality by pictures of the stimulus out of the fish’s point of view (with 360° azimuth x 180° elevation) we used a panoramic equirectangular camera (e.g. Figure 1f-h). The symbolic representations for illustrations were made as screen snapshots (e.g., Supplementary Figure S1),

Only in special tasks, such as the visualization of the light beam path and the analysis of dispersion, images were rendered with an orthographic camera in LuxCoreRender v2.1 (https://luxcorerender.org/download/) using bidirectional path tracing and appropriate volume scattering and camera clipping (e.g., Supplementary Figure S1, S2). ^30,31^.

#### Physical background

|  |  |  |
| --- | --- | --- |
| Snell's Law | $n_i \sin \varepsilon_i = n_t \sin \varepsilon_t$ | the angle of incidence ($\varepsilon_i$) and the angle of refraction/transmission ($\varepsilon_t$) with given indices of refraction $n_i$ and $n_t$ for two media. |
| Fresnel Equations | $R_s = \left\vert \frac{n_i \cos \varepsilon_i - n_t \cos \varepsilon_t}{n_i \cos \varepsilon_i + n_t \cos \varepsilon_t} \right\vert ^2$ | $R_s$, the reflectance for s-polarized light |
| Fresnel Equations | $R_p = \left\vert \frac{n_i \cos \varepsilon_t - n_t \cos \varepsilon_i}{n_i \cos \varepsilon_t + n_t \cos \varepsilon_i} \right\vert ^2$ | $R_p$, the reflectance for p-polarized light |
| | $R_{\text{eff}} = \frac{1}{2} (R_s + R_p)$ | $R_{\text{eff}}$, the total reflectance of unpolarized light |
| | $T = 1 - R$ | $T$, the fraction of the transmitted power; $R$, the fraction of the reflected power; |

#### Limits of the 3D modeling technique

Because of the limitation of our 3D modeling approach, some physical properties of the stimulation setup have not been tested in our simulations. These properties include light polarization, light interference, and diffraction. Refraction and reflection can change the polarization of light, and the intensity of the reflected light furthermore depends on the polarization of the incident light. Light interference should be absent in our experimental setup or only cause stimulus artifacts at very small spatial scales. Diffraction does not occur for the container shapes in our setup, which is in the macroscopic range well above the wavelength of visible light. Therefore, light interference and diffraction effects are not relevant for the experiments in question here.

### Animal care and transgenic lines

All animal procedures conformed to the institutional guidelines of the Universities of Tübingen and the local government (Regierungspräsidium Tübingen). The transgenic zebrafish lines *Tg(HuC:GCaMP5G)a4598Tg* was used in this study. Transgenic lines were kept in either a TL or TLN (nacre) background. Zebrafish larvae were raised in E3 medium until day 5 or 6 post-fertilization (dpf).

### Experiment protocols and data analysis

We performed the monocular direction selectivity and monocular RF mapping experiment as previously described ^8,28^, except that we used a Petri dish as a container instead of the glass bulb in the direction selectivity experiments. The data shown in the bottom of the Figure 4b have been published before as the bottom part of the Figure 1e in ^28^.

In this study, glass bulbs with diameters of 8 cm or 10 cm were used. In case of the 10 cm diameter bulb, the water level was only 1 cm above the glass bulb center. Data analysis was performed with published Matlab scripts (MOM_Load, Midbrain_Localizer and Cell_Viewer), available online (https://gin.g-node.org/Arrenberg_Lab) ^8,28,29,32^.

### Quantification and statistical analysis

The statistical information calculated with Matlab R2014b built-in functions is provided in each of the sections above. For statements of significance an alpha level of 0.05 (two-tailed) was used unless stated otherwise.

The analyzed number of zebrafish and brains is indicated in the main text and Figure legends. Error bars correspond to SEM unless stated otherwise.

## Author contributions

KW performed the experiments on tectal somatic responses. BA performed the computer graphics simulations for the three containers and the various stimulation setups. KW and JH analyzed the data. ABA, BA, KW, and JH conceived the experiments and associated analysis protocols. KW, BA, JH and ABA wrote the manuscript.

## Acknowledgments

We thank Väinö Haikala and Dierk F. Reiff for help with the visual stimulus arena. We furthermore thank Thomas Nieß (glassblower shop, University of Tübingen) and Klaus Vollmer (fine mechanics workshop, University Clinic Tübingen) for technical support. We thank David Bucciarelli and Simon Wendsche from LuxCoreRender.org for help with many questions about LuxCoreRender. This work was funded by the Deutsche Forschungsgemeinschaft (DFG) grants EXC307 (CIN – Werner Reichardt Centre for Integrative Neuroscience) and INST 37/967-1 FUGG, and a Human Frontier Science Program (HFSP) Young Investigator Grant RGY0079.

## Declaration of Interests

The authors declare no competing interests.

## Data and code availability

All raw and processed data and custom-written Matlab software used to generate the Figures will be made available upon request.

## Supplementary Figures

**Figure S1.**
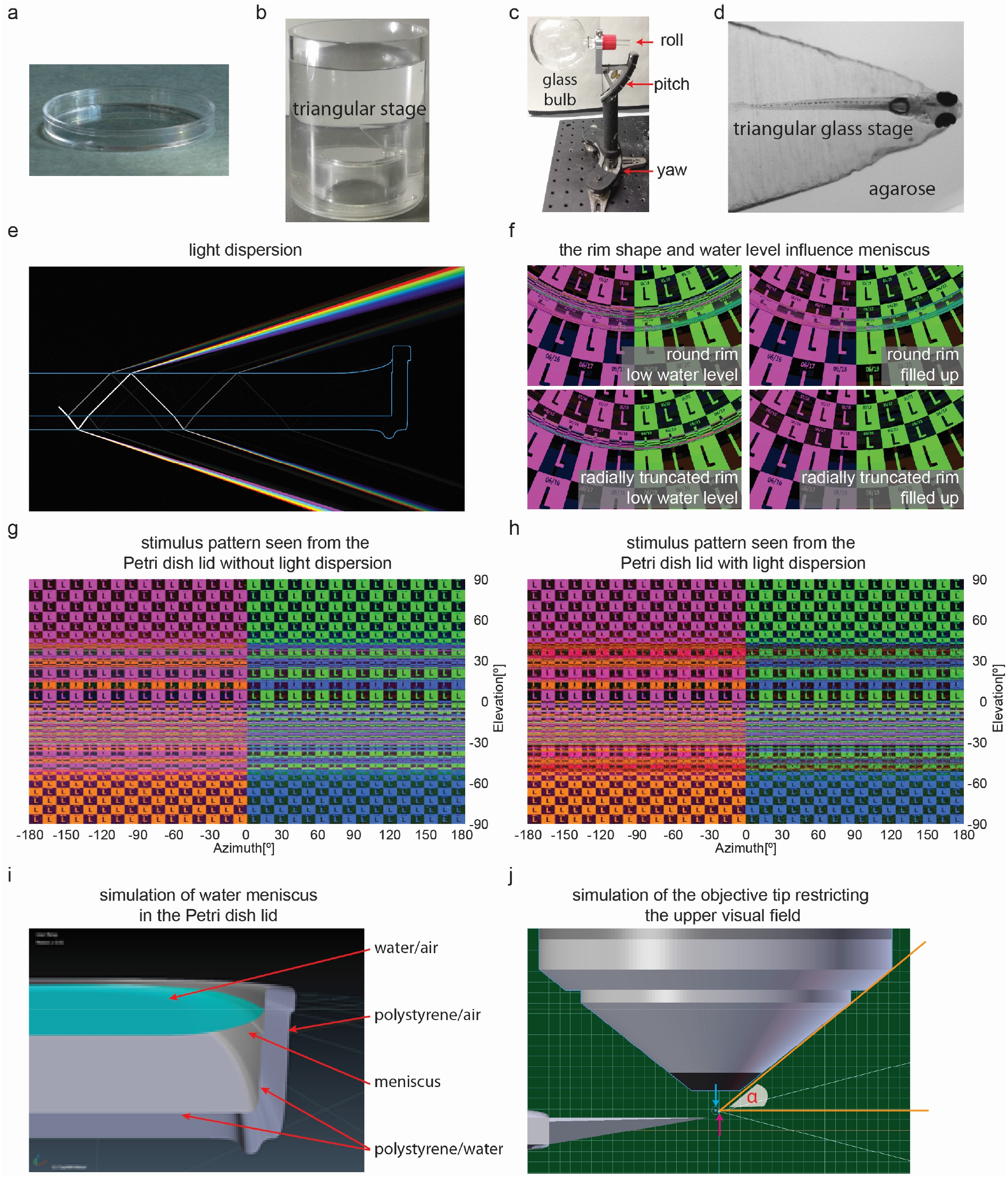
Three different containers for fish visual research and simulation of geometrical optics. Related to Figure 1. (**a**) A commercially available Petri dish lid (Ø 35 mm). (**b**) A plastic cylindrical container made by a fine mechanics workshop (Ø 40 mm). (**c**) A custom-made glass bulb (Ø 80 mm) attached to its metal holder which allows for rotational adjustment (roll, pitch, and yaw) around three axes. (**d**) A 5 dpf larval zebrafish is embedded in low melting agarose on a triangular stage in the glass bulb. (**e**) A white light beam is shed 45° downwards in the center of a Petri dish lid (side view). Because of different indices of refraction from the white light components, light dispersion occurs at the water-air and plastic-air interfaces. (**f**) The rim shape and water level influence the water meniscus in the glass bulb. Using radial truncated rim and filling up the glass bulb with water reduce the optical influences of water meniscus. The viewing perspective is from the center of the glass bulb to the stimulus point at 0° in azimuth and 50° in elevation. (**g**) The checkerboard stimulus seen from the center of the Petri dish lid (without the holder stage) without light dispersion. (**h**) The checkerboard stimulus seen from the center of the Petri dish lid (without the holder stage) with light dispersion. Color distortions and blurring exist in comparison to panel (g) but their influences are weaker than those of light infraction and reflection. (**i**) A symbolic simulation of water meniscus in the Petri dish lid (side view). On the inner wall of the Petri dish lid, the water level is higher than more central regions. (**j**) The accessible angle α (orange angle) of the fish to the visual stimulus is only 1° larger than the angle of the objective tip since the location of the fish’s eyes (magenta arrow) are roughly 0.3mm in front of the focus point (cyan arrow) of the objective.

**Figure S2.**
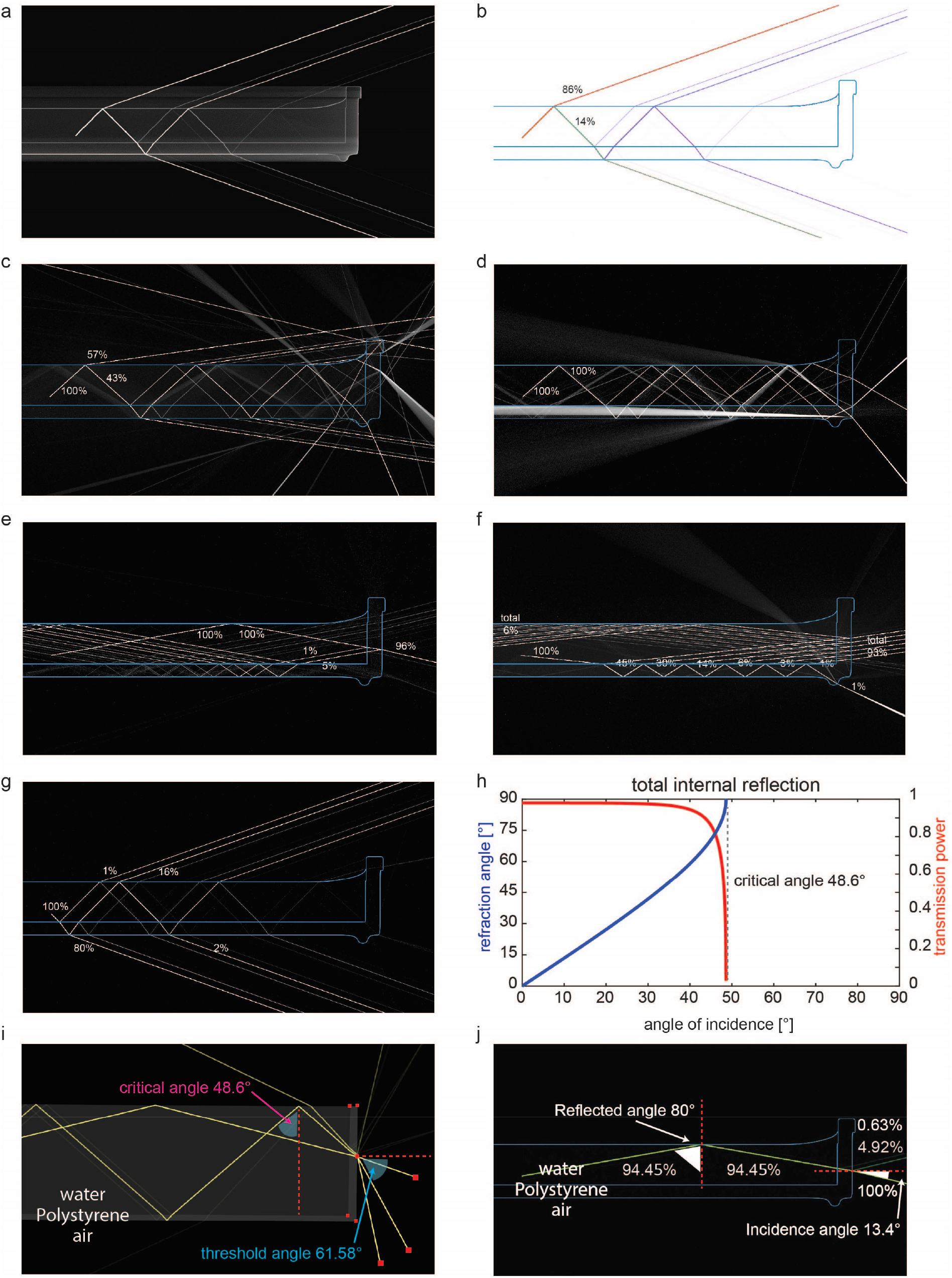
Simulation of the optics underlying visual stimulus distortion and blurring resulting from light fraction and reflection in the Petri dish lid. Related to Figure 1. For each panel from (a) to (g), only one single light beam is shown under different conditions (angles of incidence and directions). The beam splits in several rays. Because of the reversibility of the light path, this shows what the fish will see when looking in the same direction as the beam. The split rays indicate directions from which the fish will see stimulus parts. The several parts will mix to the picture for the fish with the given power of light. (**a**) A light beam is shed 45° (applicable from 0° to 48.6°) upwards. 86% of the light is refracted into the air and only 14% is reflected back. Light reflections and refractions continue to the Petri dish side wall. (**b**) The same as in panel (A) except that the light intensity is color coded (red to purple representing 100% to 2% of original light power). (**c**) A light beam is shed 48° (applicable from 0° to 48.6°) upwards. 57% of the light is refracted into the air and already 43% is reflected back. Light reflections and refractions continue to the Petri dish side wall. (**d**) A light beam is shed 49° (larger than critical angles, 48.6° from water to air and 39.6° from polystyrene to air) upwards. The light beams cannot be refracted from water or polystyrene to air. (**e**) The same as in panel (d) except that the light beam is shed 80° upwards. The light beam can only come out of the Petri dish lid through the vertical lid wall. (**f**) A light beam is shed 84° downwards. (**g**) A light beam is shed 45° downwards. (**h**) Refraction angle (in blue) and transmission power (in red) change when a light beam is projected from water (IOR = 1.333) to air (IOR = 1) and the angle of incidence increases from 0° to 90°. The critical angle of total internal reflection from water to air is indicated with a vertical dashed line. (**i**) TIR occurs on the water-air interface of the Petri dish lid when a light ray comes from below with an incidence angle smaller than 61.58°. The reflected light can reach the fish’s eye depending on the point of incidence and incidence angle. (**j**) An example light ray from panel (i). When a light ray enters the Petri dish lid with an incidence angle of 13.4° from below, 4.92% and 0.63% total light energy will be reflected on the outer and inner wall of the lid, while a light ray with 94.45% incidence light energy is totally reflected on the water-air interface.

**Figure S3.**
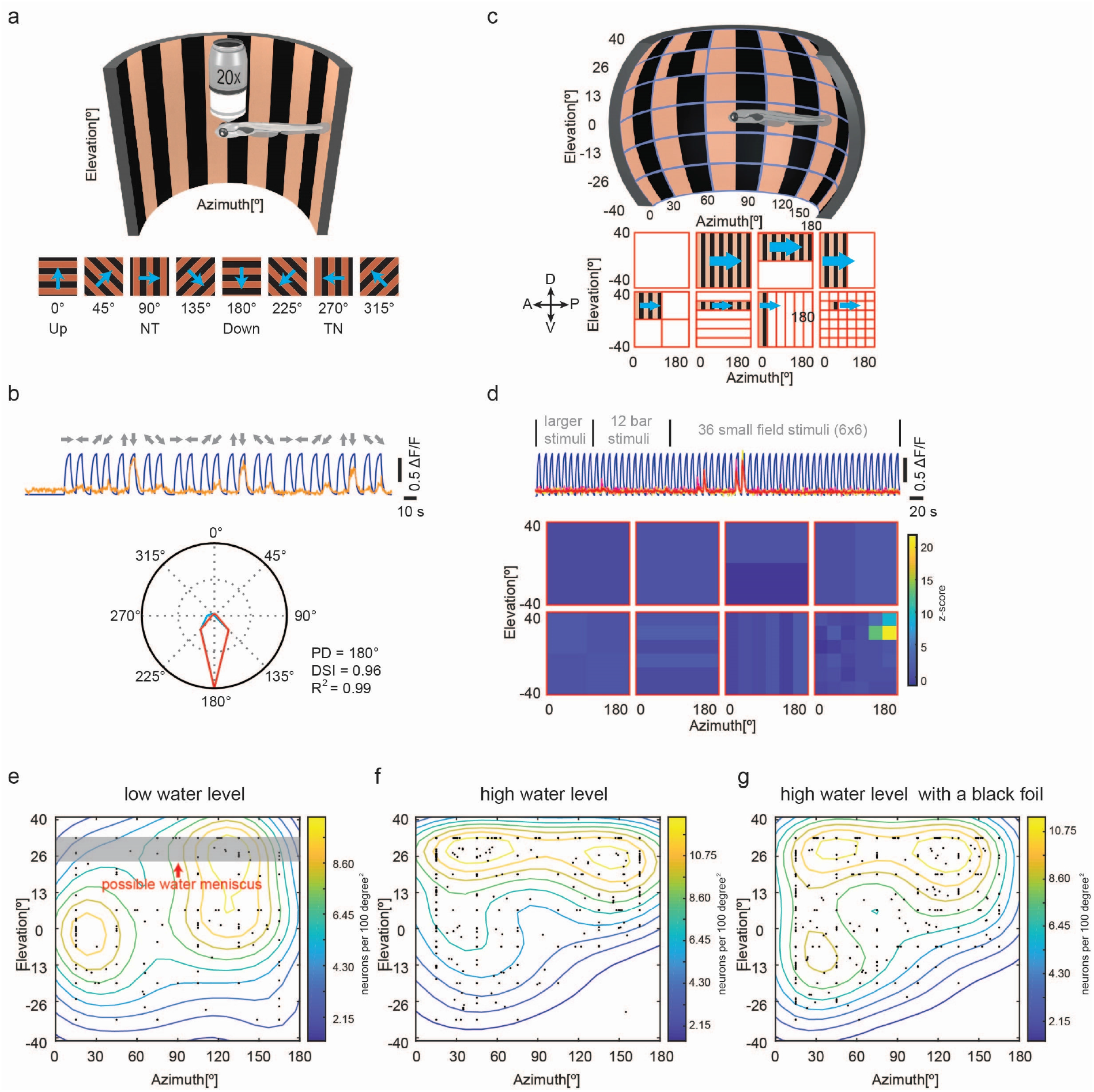
Experimental protocols for monocular direction selectivity analysis and RF mapping; low water level disrupts the distribution of receptive field centers from small-size RF tectal neurons measured with the glass bulb. Related to Figures 2, 3 and 4. (**a**) A diagram of the monocular direction selectivity analysis setup. Upper, gratings, 0.033 cycles per degree, moving in eight different directions, were presented to the right eye of the animal, which was embedded in low melting agarose in the center of the cylindrical half arena. The sizes of the fish and the arena are not proportional to the real experiments in this illustration. Below, an illustration of the 8 moving patterns. The cyan arrows indicate the moving directions. NT, nasal-temporal; TN, temporal-nasal. (**b**) An example direction-selective (DS) tectal neuron. Upper, the original calcium trace of the DS neuron in three repetitions (in orange). The blue background curve indicates the convolved (τ = 1.5 s) motion-stationary phase regressor. Below, the DS tuning curve. The neuron responded most robustly to the downward motion. (**c**) A diagram of the monocular RF mapping protocol. Upper, vertical gratings (0.033 cycles per degree) with different sizes and locations were presented to the right eye of the animal, which was embedded in low melting agarose in the center of the cylindrical half arena. The sizes of the fish and the arena are not proportional to the real experiments in this illustration. Below, an illustration of the whole visual stimulus protocol. The cyan arrows indicate the moving directions. A, anterior; P, posterior; D, dorsal; V, ventral. (**d**) The response profile of a small-size RF neuron plotted z-score. Upper, the original calcium traces (yellow, cyan and magenta) of the neuron in the three repetitions and their median (orange). The blue background curve indicates the convolved (τ = 1.5 s) motion-stationary phase regressor. Below, the response profile of a small-size RF neuron plotted with z-score. The response in each phase is corresponding to one of the motion phases in panel (c). (**e-g**) Visual field locations and density contour plot of receptive field centers of small-size RF tectal neurons recorded with low water level (panel (e), n = 4 fish, 2 composite tecta), high water level without left eye covered (panel (f), n = 6 fish, 3 composite tecta) and high water level with left eye blocked by a black foil (panel (g), n = 6 fish, 3 composite tecta), respectively. In penal (e), the low neuron density in the upper visual field potentially resulted from water meniscus (gray shade).

**Figure S4.**
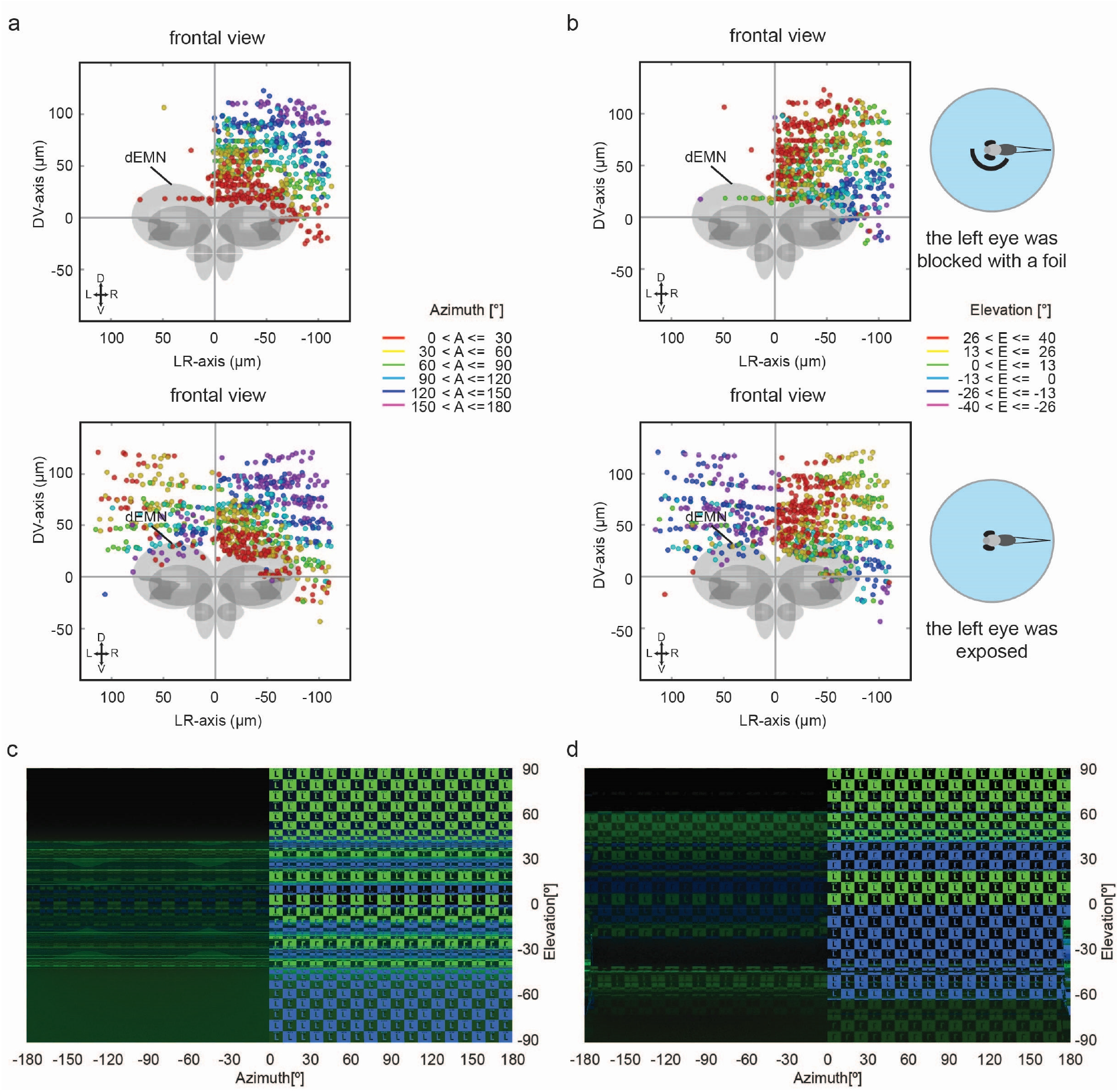
Reflection in inner surface of the glass bulb activated tectal neurons corresponding to the unstimulated eye. Related to Figure 3. (**a, b**) Frontal views of the topographic maps of tectal small-size RF neurons in azimuth (a) or elevation (b). Each colored dot represents a single neuron with its receptive field center in the corresponding azimuth (a) or elevation (b) range. For example, all receptive field centers of the neurons in red are located between 0° azimuth (in front of the fish) and 30° azimuth on the nasal right side of the fish (a). For example, receptive field centers of the neurons in green are located slightly above the equator of the view filed (0° to 13° in elevation) (b). As indicated in the illustrations to the right (dorsal view), the left eye was covered by a black foil in the top row as the control group, while the left eye was exposed to potential stimulus artifacts in the bottom row (experimental animals). n = 6 fish, 3 composite brains. (**c, d**) Light reflection artifacts for the Petri dish lid (c) and the cylindrical container (d) using a monocular stimulus (cf. Figure 3b).

**Figure S5.**
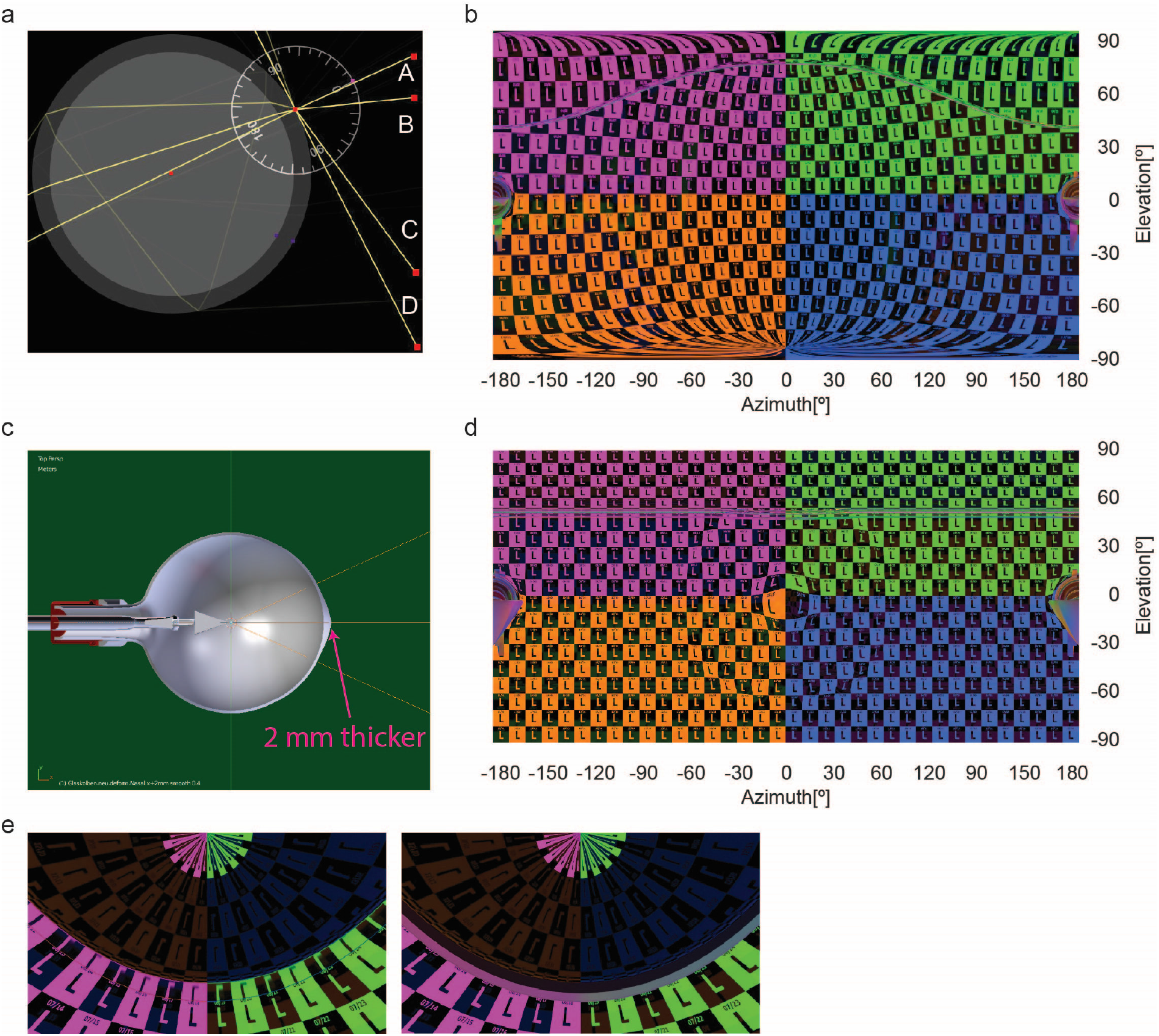
Other factors which influence the visual stimulus patterns perceived by the fish. Related to the discussion. (**a, b**). (**a**) Four light beams (A, B, C and D) are shed into the glass bulb with different angles of incidence simulated with software (Ray Optics Simulation). The indices of refraction from water (inner circle) and glass (outer ring) are 1.333 and 1.47. The reflected light reaches the unstimulated eye (close to the center of the glass bulb) of the fish (left eye) only when the incidence angle of the visual stimulus is small (e.g., beam A). (**b**) An ideal checkerboard stimulus perceived when the animal is located far in front of the glass bulb center (15 mm = 38% of the radius). (**c**) Dorsal view of a simulated glass bulb with the frontal point 200% thicker (3 mm instead of 1 mm) than other regions of the glass wall. (**d**) An ideal checkerboard stimulus perceived by a fish when the animal is located in the center of the glass bulb shown in (c). (**e**) The material around the objective lens can cause additional stimulus reflections and it is advantageous to use objectives with non-reflective surfaces. Reflection of the stimulus pattern by microscope objective tips with different textures. Left side, glossy metal results in upside down reflection (upside down letters ‘L’); right side, Matt black and white, like Zeiss W N-Acroplan.

## Notes

### Competing Interest Statement

The authors have declared no competing interest.

